# GENERator-v2: Reconciling Coarse Tokenization with Single-Nucleotide Resolution in Genomic Language Modeling

**DOI:** 10.64898/2026.01.27.702015

**Authors:** Qiuyi Li, Zhihao Zhan, Shikun Feng, Yiheng Zhu, Yuan He, Wei Wu, Zhenghang Shi, Shengjie Wang, Zongyong Hu, Zhao Yang, Jiaoyang Li, Jian Tang, Haiguang Liu, Tao Qin

## Abstract

Genomic foundation models aim to learn general-purpose representations directly from DNA sequence, enabling sequence understanding, generation, and probabilistic reasoning across a wide range of biological tasks. Scaling such models to genomic lengths, however, remains challenging due to the tension between long-range context, nucleotide-level resolution, and practical computational efficiency. Architectural innovations have enabled increasingly long nominal inputs, but often struggle to translate additional context into meaningful performance gains, particularly in the presence of sparse functional signal along eukaryotic genomes. In this work, we revisit the design of long-context genomic foundation models from the perspective of training objective and data construction. We introduce Factorized Nucleotide Supervision (FNS), which reconciles efficient *k*-mer tokenization with single-nucleotide likelihoods through probability marginalization, and Genome Compression Pretraining (GCP), which reshapes the training distribution by concentrating on gene-centric and regulatory regions. Together, these techniques enable standard transformer-based models to perform functional in-context learning without sacrificing nucleotide-level fidelity or computational efficiency. Building on these ideas, we present a family of autoregressive genomic foundation models supporting contexts of up to 98k base pairs across eukaryotic and prokaryotic genomes. Across training-free evaluations and downstream fine-tuning benchmarks, our models consistently improve over prior approaches and match or exceed state-of-the-art baselines while enabling substantially more efficient inference. Together, these results demonstrate that aligning supervision and data regimes with the biological structure of genomic sequence provides a principled and effective path toward scalable and biologically faithful genomic language modeling. Models, data, and scripts for downstream analyses are publicly available at https://huggingface.co/GenerTeam.

## Introduction

The rapid growth of high-throughput sequencing technologies has led to an unprecedented accumulation of genomic data across a wide range of organisms [29]. This expansion of available sequence data has motivated increasing interest in genomic foundation models (GFMs) [40], which aim to learn general-purpose representations directly from raw DNA sequence. Such models are designed to support a wide range of downstream tasks, including genome annotation, variant interpretation, and sequence design. Inspired by large language models in natural language processing [38], recent work has explored a variety of self-supervised learning paradigms for modeling genomic sequences at scale.

Existing GFMs can be broadly categorized by their training objective and model architecture. Masked language models (MLMs, [11]), such as DNABERT-2 [42], the Nucleotide Transformer (NT) [9], dominate early efforts in genomic representation learning. These models demonstrate strong performance on sequence comprehension and discriminative prediction tasks, but their bidirectional masking objective limits their generative capability and makes them less suitable for open-ended sequence generation and probabilistic reasoning.

Autoregressive models [1] address this limitation by modeling the joint distribution of genomic sequence via next-token prediction. Representative examples include HyenaDNA [28], GENERator [41], and Evo2 [7]. More recently, discrete diffusion-based generative models have also been explored for genomic sequence modeling [35, 33, 2, 10], providing an alternative framework to autoregressive generation. Despite their differences, all of these approaches face a common challenge when scaling to genomic lengths: balancing long-range context, nucleotide-level resolution, and computational efficiency.

Long-context modeling is particularly important in genomics, as meaningful genomic prediction often requires conditioning over multiple gene-centric and regulatory units distributed across large genomic intervals [26, 12, 20]. Genomic sequences are most naturally represented at single-nucleotide resolution, and many genomic foundation models adopt single-nucleotide tokenization as their base representation. While this choice preserves biological fidelity, modeling long genomic context at single-nucleotide resolution is computationally prohibitive for standard Transformer architectures, giving rise to a fundamental tension between long-context modeling, single-nucleotide fidelity, and practical computational budgets.

A variety of architectural strategies have been proposed to mitigate this tension under single-nucleotide tokenization. Several models incorporate convolutional components to extend effective context length, including Enformer [3], HyenaDNA [28], Evo [27], Evo2 [7], and AlphaGenome [4], which augment sequence modeling with Conv1D or convolution-like operators to increase receptive fields. In parallel, state-space models (SSMs), such as Caduceus [37] and HybriDNA [23], replace explicit attention with recurrent-style hidden state updates that scale linearly with sequence length. While these approaches support much longer nominal inputs, prior studies have shown that their ability to exploit additional context often saturates as sequence length increases [41]. As context grows, increasingly large portions of the sequence must be compressed into fixed-size hidden representations, limiting effective long-range understanding.

An alternative approach addresses the same challenge by reducing the number of tokens processed by the model. Tokenization-based compression allows each token to represent multiple nucleotides, thereby extending the effective genomic span accessible to Transformer architectures. Broadly, such approaches fall into two categories. The first category comprises learnable tokenization schemes. This includes explicit subword tokenization based on byte-pair encoding (BPE, [16]), adopted by models including GROVER [34], DNABERT-2 [42], and GenomeOcean [43], as well as *token-merging* approaches such as VQDNA [18], MxDNA [31], and mergeDNA [19], which learn to aggregate nucleotides into higher-level units during training [6]. While effective in masked language modeling, these approaches introduce intrinsic ambiguity in autoregressive generation: overlapping or hierarchical tokens lead to multiple plausible next-token predictions, resulting in unstable training and degraded generative performance [41]. The second category is fixed-length *k*-mer tokenization, which represents genomic sequence as non-overlapping blocks of length *k*. This strategy, adopted by DNABERT [15], the Nucleotide Transformer [9], and GENERator [41], avoids hierarchical ambiguity and provides a well-defined autoregressive objective. Nevertheless, *k*-mer tokenization has been widely criticized for discarding single-nucleotide resolution and for its sensitivity to tokenization phase.

Among existing autoregressive models, Evo2 [7] represents a notable advance by combining single-nucleotide tokenization with a StripedHyena architecture that embeds convolutional structure into the model [7]. While this design is more efficient than a vanilla Transformer, it still incurs substantial computational cost, and its ability to convert additional context into meaningful performance gains remains limited in practice. These observations suggest that purely architectural solutions may not fully resolve the fundamental trade-offs between context length, resolution, and efficiency.

In this work, we pursue a different approach. Rather than entangling architectural design with additional complexity, we focus on aligning the training objective and data distribution with the biological structure of genomic sequence.

We retain the Transformer architecture and *k*-mer tokenization, but revisit genomic language modeling along the dimension of training objective. First, we introduce *Factorized Nucleotide Supervision (FNS)*, which decomposes each *k*-mer prediction into *k* single-nucleotide likelihoods via marginalization, …

We retain the Transformer architecture and *k*-mer tokenization, but address their perceived limitations through two complementary techniques. First, we introduce *Factorized Nucleotide Supervision (FNS)*, which decomposes each *k*-mer prediction into *k* single-nucleotide likelihoods via marginalization, thereby restoring single-nucleotide resolution without sacrificing the efficiency of coarse tokenization. This objective bears conceptual similarity to multi-token prediction strategies used in large language models [22], but operates in the reverse direction: instead of predicting multiple tokens per step, it factorizes a single token into multiple biologically meaningful components.

Second, we propose *Genome Compression Pretraining (GCP)*, a data construction strategy designed to address the extreme sparsity of functional signal in eukaryotic genomes. By concatenating gene-centric and regulatory regions while discarding large stretches of low-information background, GCP enables effective utilization of functional context and induces an explicit next-gene prediction signal. This design implements, at the data level, an effect analogous to sparse attention, while preserving the stability and simplicity of dense self-attention.

Building on these ideas, we introduce *GENERator-v2*, a family of autoregressive genomic foundation models trained with FNS and GCP. GENERator-v2 supports genomic contexts of up to 98k base pairs and includes domain-specialized variants for both eukaryotic and prokaryotic genomes, enabling modeling across all domains of life. Across extensive training-free evaluations and downstream fine-tuning on revised NT tasks, GENERator-v2 demonstrates consistent improvements over the original GENERator model [41]. Moreover, GENERator-v2 achieves performance that matches or exceeds Evo2 on key generative and probabilistic tasks, while being substantially more efficient in inference.

Collectively, these results suggest that progress in scalable genomic modeling requires more than architectural innovation alone. In particular, our findings highlight the importance of aligning supervision and data regimes with the biological structure of genomic sequence in order to convert increased context length and model capacity into meaningful performance gains.

## 2 Factorized Nucleotide Supervision (FNS)

### 2.1 Motivation

Non-overlapping *k*-mer tokenization offers substantial computational advantages for long-context genomic language modeling. By reducing sequence length by a factor of *k*, it enables efficient scaling to contexts on the order of 10^5^ nucleotides. However, treating each *k*-mer as an atomic prediction target introduces a systematic limitation at the level of the training objective.

Under standard next-token cross-entropy, a model trained with a *k*-mer vocabulary of size 4^*k*^ receives exactly one correct label at each step. For *k* = 6, this means selecting one class among 4096 possibilities. All incorrect *k*-mers are penalized equally, regardless of their nucleotide-level similarity to the ground truth. Consequently, predictions that differ by a single nucleotide and those that are entirely mismatched incur identical loss.

This supervision is maximally sparse and biologically misaligned. Genomic sequences exhibit high background entropy and frequent near-miss variants (e.g., SNPs), where graded similarity is meaningful. As training progresses, the model must resolve fine-grained distinctions that are invisible under discrete *k*-mer supervision, often leading to unstable late-stage optimization and poor calibration under single-nucleotide perturbations.

### 2.2 Method Overview

Factorized Nucleotide Supervision (FNS) addresses these objective-induced issues by redefining the learning signal while preserving the computational structure of *k*-mer tokenization. The core idea is to reinterpret each *k*-mer token as a structured object composed of *k* nucleotides. Instead of supervising the model with a single categorical target over 4^*k*^ classes, FNS derives nucleotide-level conditional probabilities from the same logits and applies supervision independently at each nucleotide position.

Notably, FNS does not modify the model architecture, tokenizer, or output dimensionality. The model still produces 4^*k*^ logits per step; FNS operates entirely at the level of loss construction and gradient allocation. An overview of the difference between naive *k*-mer supervision and FNS is shown in Figure 1. Formal definitions, theoretical results, and algorithmic details are provided in Appendix A.

**Figure 1:**
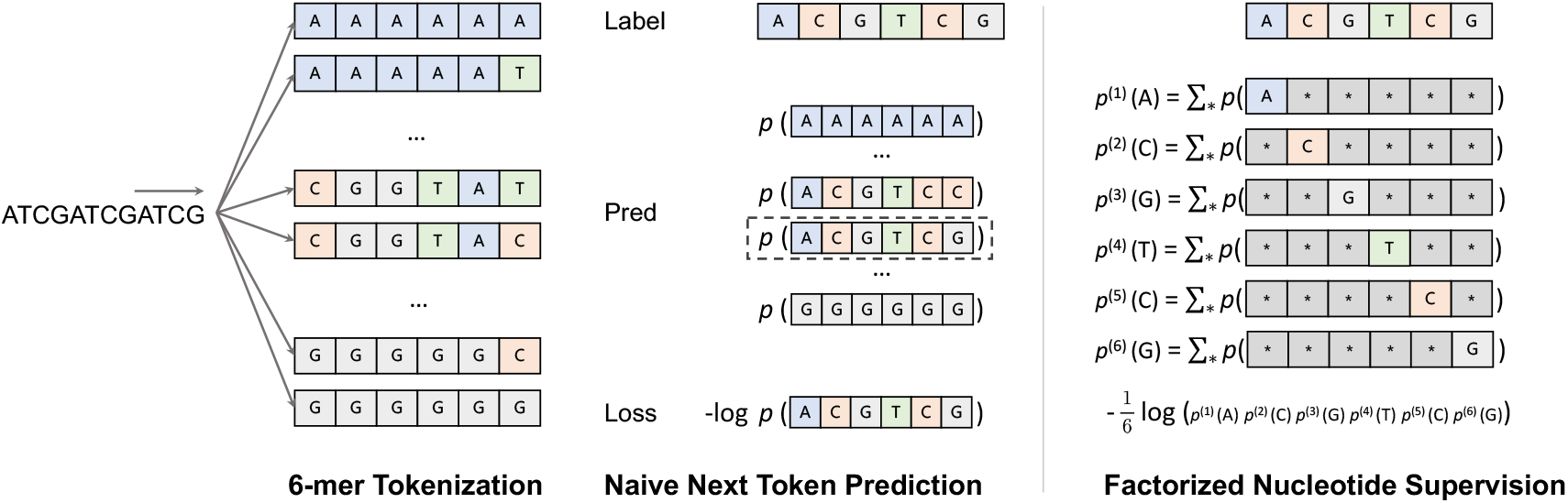
Illustration of Factorized Nucleotide Supervision (FNS). Non-overlapping *k*-mer tokenization maps a nucleotide sequence into a discrete token space of size 4^*k*^, where many tokens differ by only a small number of nucleotides. Under naive next-token prediction, the model assigns probability mass to entire *k*-mer tokens and is supervised only on the exact target token, such that partially correct predictions incur the same loss as completely incorrect ones. FNS addresses this limitation by projecting token-level probabilities onto nucleotide-level marginals via summation over all tokens consistent with each nucleotide at each position. The training loss is then computed as the sum of per-nucleotide negative log-likelihoods, yielding dense, graded supervision while preserving the original *k*-mer output space and model architecture.

### 2.3 Formulation

Let 𝒱 = ℬ^*k*^ denote the *k*-mer vocabulary, where ℬ = {*A, C, G, T*}. Given an input context, the model outputs logits **z** ∈ ℝ^|ℬ|^, inducing a categorical distribution *p*(*v*) = softmax(**z**)_*v*_ over *k*-mers. For each nucleotide position *j* ∈ {1, …, *k*} and base *b* ∈ ℬ, FNS defines the induced nucleotide marginal

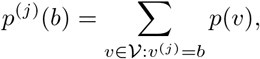

where *v*^(*j*)^ is the *j*-th nucleotide of *v*. Given a ground-truth *k*-mer target *y* = (*y*^(1)^, …, *y*^(*k*)^), the FNS loss is

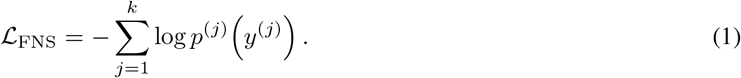

This construction yields nucleotide-level supervision while remaining fully differentiable with respect to the original 4^*k*^ logits, such that partially correct predictions receive proportionally smaller penalties than completely incorrect ones.

### 2.4 Practical Implications

FNS admits a clean probabilistic interpretation: training with next *k*-mer prediction under FNS is equivalent to maximum-likelihood training of a predictor that outputs *k* single-nucleotide tokens per step under a conditional-independence assumption given the shared hidden state (formal statement and proof in Appendix A.2). Operationally, this interpretation explains two practical benefits.

#### Optimization stability

Under standard *k*-mer cross-entropy, gradient signal is dominated by the single target class. Under FNS, all logits contribute via nucleotide marginals, yielding denser gradients and a smoother optimization landscape, particularly in late training stages.

#### SNP-level reasoning

Because FNS produces nucleotide-level conditional probabilities, the model can score single-nucleotide variants via likelihood ratios without retraining or architectural modification (derivation in Appendix A.2). This capability is critical for variant effect prediction and other SNP-sensitive downstream tasks.

### 2.5 Limitations and Summary

FNS assumes conditional independence among nucleotides within a *k*-mer given the hidden state. While this assumption is not strictly true for DNA, it is a controlled approximation that trades a small modeling bias for substantial gains in stability, resolution, and data efficiency. Dependencies within *k*-mers can still be captured implicitly through the shared hidden representation.

In summary, FNS redefines *k*-mer language modeling as a structured, nucleotide-aware learning problem. By decomposing discrete *k*-mer prediction into marginal nucleotide supervision, it alleviates label sparsity, stabilizes optimization, and enables SNP-level reasoning while preserving the computational advantages of coarse tokenization.

## 3 Genome Compression Pretraining (GCP)

### 3.1 Motivation

Genome Compression Pretraining (GCP) is motivated by a fundamental mismatch between the distribution of functional signal in eukaryotic genomes and the fixed-length context windows used in modern long-context genomic foundation models.

Early genomic foundation models, such as DNABERT [15] and the Nucleotide Transformer [9], primarily adopted an *all-sequence training* paradigm, in which entire genomes were directly segmented and used for self-supervised learning. While straightforward, this approach expends the majority of model capacity on weakly constrained or low-information background sequence, reflecting the extreme sparsity of functional signal in eukaryotic genomes.

GENERator was among the first models to explicitly address this issue through *functional sequence training*, which restricts pretraining to annotated functional regions. Extensive evaluation demonstrated that concentrating training on gene-centric and regulatory regions substantially improves representation quality and downstream performance. Subsequent large-scale models, including Evo2 [7] and Genos [21], converged on similar insights and also incorporated annotation-aware data selection strategies.

However, functional sequence training introduces a new limitation when combined with long-context autoregressive models. Most individual functional regions are far shorter than the nominal context length supported by modern architectures. As a result, training rarely requires conditioning across multiple distant functional units, leading to systematic under-utilization of long-context capacity and limited modeling of interactions among functional regions.

One approach to mitigating this issue, adopted by Evo2, is to decouple learning local functional structure from long-range genomic context. In practice, this involves pretraining with relatively short contexts (e.g., 8k base pairs) on functional regions, followed by large-scale context expansion to megabase-scale genomic segments that include both functional sequence and background. While effective, this strategy relies on reintroducing large amounts of background sequence and incurs extremely high computational cost, requiring substantial specialized hardware to train and scale.

GCP adopts a different perspective. Rather than expanding context by reintroducing large amounts of background sequence, GCP reshapes the training data distribution itself. By concatenating gene-centric and regulatory regions in genomic order, GCP enables long-context models to operate on densely informative sequences, encouraging explicit modeling of interactions among functional units while preserving computational efficiency.

### 3.2 Method Overview

GCP is a data-level pretraining strategy designed to align long-context modeling with the functional organization of eukaryotic genomes. Rather than modifying model architecture or attention mechanisms, GCP reshapes the training data distribution so that standard dense self-attention operates over sequences that are densely populated with biologically informative content. As a result, long-context capacity is devoted primarily to modeling relationships among functional genomic units, rather than expended on low-information background sequence.

Concretely, GCP constructs a *compressed genome* by selecting annotated gene-centric and regulatory regions, ordering them by genomic coordinate, and concatenating the corresponding sequences into a single stream with separator tokens (<s>) marking boundaries between adjacent regions. Separator tokens are included in the input but masked from the language-modeling loss. Fixed-length training windows are then sampled from this compressed stream. Because background sequence has been removed, each window consistently saturates the available context length with functional genomic sequence.

As a result, a single training window typically contains multiple neighboring gene-centric and regulatory regions arranged in genomic order; for models such as GENERator with a 98k base-pair context, this corresponds to dozens of functional units within a single forward pass. These regions collectively form a *functional context* that conditions predictions for downstream regions on upstream functional units, enabling the model to exploit co-occurrence and neighborhood structure during next-token prediction. This behavior mirrors in-context learning in large language models and is therefore referred to as *functional in-context learning*.

A key design choice in GCP is the definition of functional regions. We adopt a broad and conservative, gene-centric definition based on RefSeq [29] annotations. Functional regions are defined at the level of annotated gene entries, rather than finer-grained units such as individual coding sequences or exons. For protein-coding genes, extracted regions span extended genomic loci that may include multiple CDS segments across exons, intervening introns, untranslated regions (UTRs), and associated regulatory sequence where annotated. The same definition applies to RNA genes. This gene-aware formulation intentionally preserves substantial contextual redundancy around genes, prioritizing information preservation and representation robustness over maximal compression.

Under this definition, GCP achieves an approximate compression ratio of 1:5 on eukaryotic genomes. While protein-coding CDS typically account for only a small fraction of the genome, the gene-centric regions retained under GCP cover roughly one-fifth of the genomic sequence. In this sense, genome compression favors information preservation: low-information background sequence is removed, while broad functional neighborhoods around genes are retained. An overview of the GCP pipeline is shown in Figure 2, with formal definitions, theoretical analyses, and algorithmic details provided in Appendix B.

**Figure 2:**
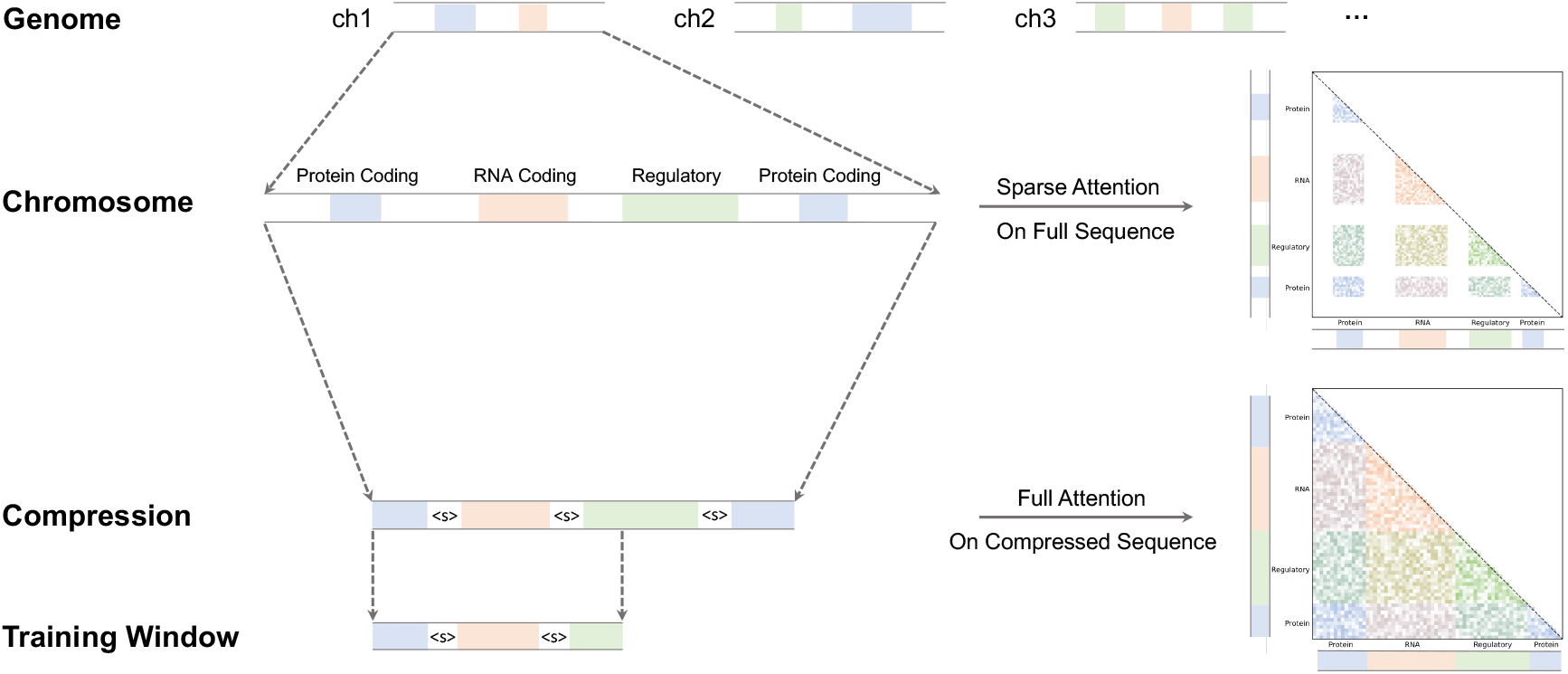
Overview of Genome Compression Pretraining (GCP). Starting from a full genome composed of multiple chromosomes, annotated functional regions—including protein-coding genes, RNA-coding genes, and regulatory elements—are first identified along each chromosome. These regions are then concatenated in genomic order to form a compressed sequence, with separator tokens (<s>) marking boundaries between adjacent units. Fixed-length training windows are sampled from the compressed stream, ensuring that each window contains a dense collection of biologically informative sequence. By reshaping the training data distribution, GCP enables standard dense self-attention to operate directly on informative regions. This data-level sparsification achieves an effect analogous to sparse attention on the full genome, while avoiding heuristic token selection and preserving stable representation learning and effective context utilization.

### 3.3 Formulation

Let a chromosome *C* be associated with a set of annotated regions 𝒜
(*C*) = {*r*_*i*_}, where each region corresponds to a gene-centric or regulatory unit. Regions are grouped into three broad categories: (i) protein-coding genes, (ii) RNA-coding genes, and (iii) regulatory regions. For protein-coding genes, each region spans an extended genomic neighborhood that includes not only CDS segments, but also introns, UTRs, promoters, and intervening non-coding sequence within the locus; gene-proximal intergenic sequence may also be included to preserve local context.

After sorting regions by genomic position, we tokenize each region independently and concatenate them using a separator token <s>:

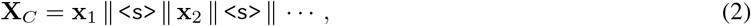

where **x**_*i*_ denotes the tokenized sequence corresponding to region *r*_*i*_. Training proceeds using standard autoregressive next-token prediction over fixed-length windows of *L* tokens sampled from the compressed sequence, with separator tokens masked from the language modeling loss.

We emphasize that while genomic *order* is preserved, genomic *distance* is not. This is intentional: GCP is designed to model neighborhood structure and co-occurrence patterns among gene-centric and regulatory units rather than raw base-pair contiguity.

### 3.4 Practical Implications

#### Context utilization and effective span

By construction, compressed genomic sequences are no longer constrained by raw base-pair contiguity. As a result, training windows sampled from the compressed stream can consistently saturate the full context length *L* with biologically informative sequence, rather than low-information background. A window of length *L* in compressed space typically spans multiple gene-centric and regulatory units in their genomic order, while often covering loci that are widely separated in base-pair distance in the original genome. In practice, this yields a substantially larger *effective conditional span*, enabling models to condition on interactions among many functional units without architectural modification.

#### Tokenization-shift augmentation under FNS

Non-overlapping *k*-mer tokenization is inherently phase-sensitive: shifting the tokenization offset by one nucleotide produces a completely different sequence of token IDs. To mitigate sensitivity to arbitrary token boundaries, we employ an exhaustive tokenization-shift augmentation strategy, cycling through all *k* possible offsets across training. Under Factorized Nucleotide Supervision (FNS), the aggregate training objective across offsets induces complete and consistent nucleotide-level supervision up to boundary effects, which can be eliminated by padding (Proposition 4). This ensures that genome compression and tokenization-shift augmentation do not introduce systematic bias at the nucleotide level.

#### Data-level sparsity versus sparse attention

Sparse attention mechanisms reduce computational cost by restricting which tokens participate in attention, but often rely on heuristic token selection and can lead to unstable or discontinuous representations. In contrast, GCP enforces sparsity at the data level: only selected functional tokens appear in a context window, but all present tokens participate fully in dense self-attention. This preserves the smoothness properties of dense attention while substantially reducing the effective sequence length.

#### Implications for representation stability

In autoregressive decoders, a common practice is to extract a sequence embedding from the hidden state of a designated token (e.g., the final token), which aggregates information from the entire context via self-attention. Under sparse attention, the representation of such a token can depend sensitively on the selected attention subset, and small changes in token selection may induce large, discontinuous changes in the resulting embedding. By contrast, because GCP performs sparsification prior to attention and applies dense self-attention over all retained tokens, the resulting embeddings vary smoothly with respect to the input sequence. This stability is particularly important for representation-centric genomic tasks, and provides an additional justification for implementing sparsity at the data level rather than within the attention mechanism itself.

### 3.5 Limitations and Summary

GCP biases pretraining toward annotated gene-centric and regulatory regions while intentionally retaining redundancy, particularly within protein-coding gene neighborhoods. This conservative compression strategy prioritizes information preservation over aggressive pruning, at the cost of relying on annotation quality and completeness.

In summary, Genome Compression Pretraining provides a simple, robust, and architecture-agnostic mechanism for long-context genomic modeling. By enforcing sparsity through data construction rather than attention heuristics, GCP improves context utilization, preserves training stability, and yields representations well-suited for downstream genomic analysis.

## 4 GENERator-v2

### 4.1 Model Overview

We introduce GENERator-v2, an autoregressiv genomic foundation model trained with revised pre-training strategies. At the architectural level, GENERator-v2 follows the original GENERator design: all variants employ the same Transformer decoder backbone, non-overlapping 6-mer tokenization, and a maximum supported context length of 98k base pairs. Unless otherwise specified, optimization settings and long-context engineering choices are also inherited from the v1 configuration.

This design enables a controlled comparison between model generations. By holding the architecture and tokenization scheme fixed, differences between GENERator-v1 and GENERator-v2 can be attributed primarily to changes in training objective and data construction, rather than to confounding architectural factors.

### 4.2 Domain-Specific Variants

GENERator-v2 is released in two domain-specialized variants, reflecting fundamental differences in functional organization between eukaryotic and prokaryotic genomes:

- GENERator-v2-eukaryote, designed for eukaryotic genomes, and
- GENERator-v2-prokaryote, designed for prokaryotic genomes.

For each domain, we train two model scales, with approximately 1B and 3B parameters, yielding four models in total. This configuration supports systematic analysis across biological domains and model scales under a unified training and evaluation protocol.

### 4.3 Pre-training Strategy

All GENERator-v2 models are trained using Factorized Nucleotide Supervision (FNS; Section 2), which replaces discrete *k*-mer supervision with nucleotide-level likelihoods induced from *k*-mer logits. Because FNS operates purely at the level of the training objective, it is applied uniformly across both eukaryotic and prokaryotic variants without architectural modification.

Genome Compression Pretraining (GCP; Section 3) is applied selectively to the eukaryotic models. This choice reflects differences in functional density between domains. Eukaryotic genomes exhibit sparse functional signal, with annotated functional regions accounting for a small fraction of the total sequence. In this setting, naïve long-context training expends substantial capacity on low-information background, whereas GCP concentrates training on gene-centric and regulatory regions to improve effective context utilization.

Prokaryotic genomes, by contrast, are highly compact, with functional sequence often comprising the majority of the genome. In this regime, aggressive compression offers limited benefit and may discard biologically informative context, particularly in the absence of comprehensive annotations. Accordingly, GCP is not applied to GENERator-v2-prokaryote.

### 4.4 Training Data

The two domain-specific variants differ in training data composition, reflecting the requirements of their respective pre-training pipelines.

#### Eukaryotic training data

GENERator-v2-eukaryote is trained exclusively on genomes from RefSeq [29], as GCP relies on high-quality gene and regulatory annotations to construct compressed training sequences. The curated nature of RefSeq provides a reliable foundation for annotation-driven pre-training.

#### Prokaryotic training data

GENERator-v2-prokaryote is trained on a combined corpus drawn from RefSeq and the Genome Taxonomy Database (GTDB) [30]. GTDB offers broader phylogenetic coverage and a more balanced distribution of prokaryotic species, complementing RefSeq data. Although GTDB lacks detailed functional annotations, this limitation is compatible with the prokaryotic setting, where dense functional coverage reduces reliance on annotation-driven preprocessing.

An overview of all GENERator variants and their training configurations is provided in Table 1.

**Table 1:** Summary of the GENERator model family. All models share the same Transformer decoder architecture, 6-mer tokenization, and 98k context length. GENERator-v2 models differ only in training data composition and pre-training strategy (FNS/GCP), enabling a controlled comparison.

| Model | Domain | Params | FNS | GCP | Data Source |
| --- | --- | --- | --- | --- | --- |
| GENERator | Eukaryote | 1.2B / 3B | ✗ | ✗ | RefSeq |
| GENERator-v2-eukaryote | Eukaryote | 1.2B / 3B | ✓ | ✓ | RefSeq |
| GENERator-v2-prokaryote | Prokaryote | 1.2B / 3B | ✓ | ✗ | RefSeq + GTDB |

## 5 Experiments

### 5.1 Sequence Recovery

We first examine the effect of Factorized Nucleotide Supervision (FNS) using *sequence recovery* as a unified, training-free evaluation protocol. This task jointly evaluates intrinsic generative accuracy and generation efficiency under identical input–output settings, enabling fair comparison across models with different tokenization schemes and architectures.

#### Evaluation protocol

Genomic sequences are grouped by taxonomic category. For each group, we randomly sample genomic segments of length 6,144 base pairs from RefSeq and use each segment as input to generate the subsequent 30 nucleotides. The generated sequence is then compared against the ground-truth sequence to compute recovery accuracy. To ensure biological relevance of the task, we impose two constraints: (i) the rightmost end of each input segment must lie within a coding sequence (CDS), and (ii) at least two-thirds of each segment must consist of non-coding sequence. The first constraint ensures a well-defined coding context for generation, while the second evaluates models in settings where long-range, non-coding context is available.

#### Results on eukaryotic genomes

On eukaryotic datasets, GENERator and Evo2 exhibit highly consistent performance patterns across all taxonomic groups. Relative to GENERator-v1, GENERator-v2 shows stable and systematic improvements, indicating the benefit of the revised pre-training strategy. At matched parameter scales, GENERator-v2-1B outperforms Evo2-1B, while GENERator-v2-3B achieves performance comparable to Evo2-7B despite the substantial difference in model size (Figure 3A). These results are consistent with the role of FNS in compensating for the coarser granularity introduced by 6-mer tokenization, enabling improved generative accuracy without sacrificing efficiency.

**Figure 3:**
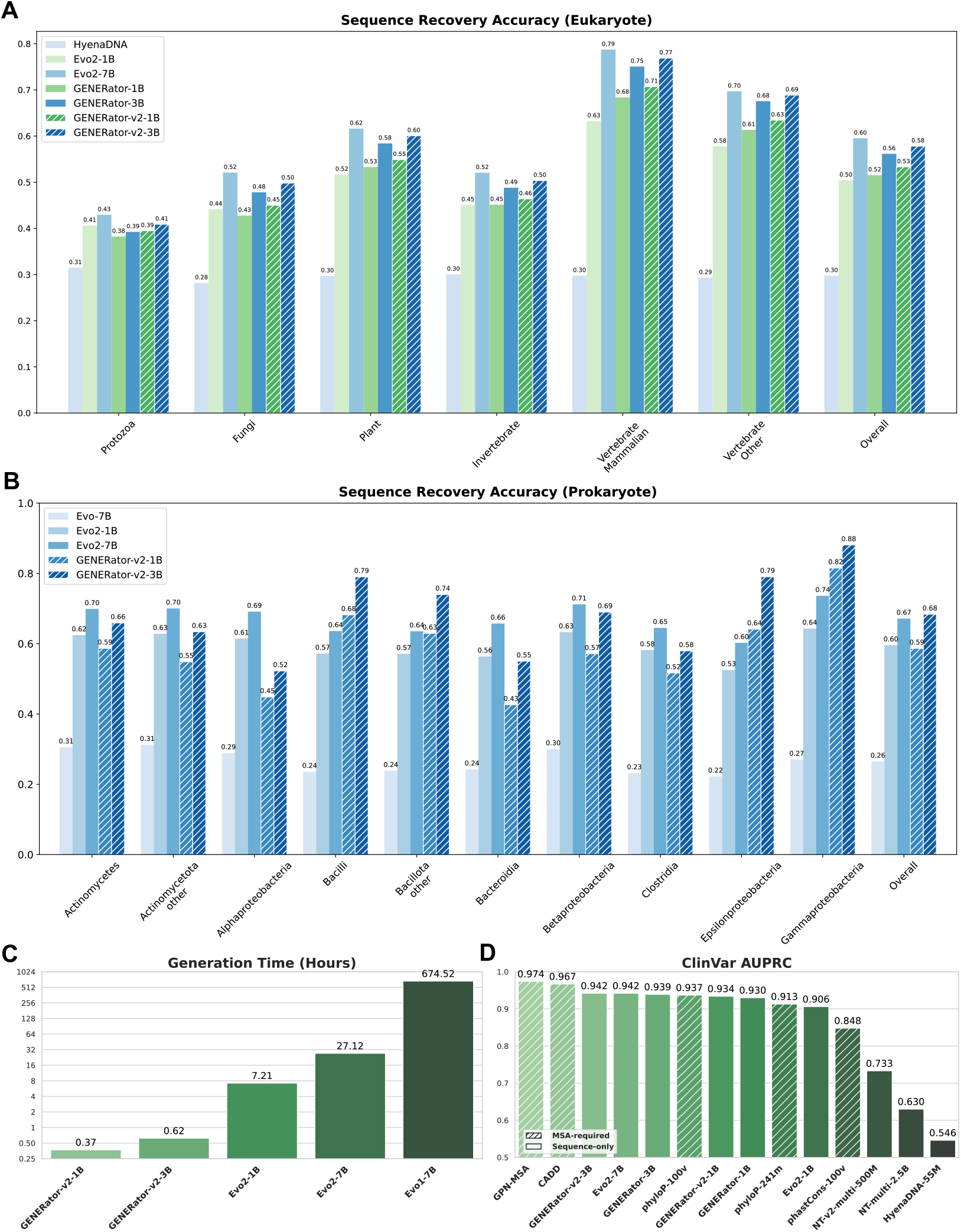
Summary of training-free evaluations across intrinsic generation accuracy, computational efficiency, and variant effect prediction. Sequence recovery accuracy is reported for eukaryotic genomes (A) and prokaryotic genomes (B), grouped by taxonomic category under a fixed input length of 6,144 base pairs and a fixed generation target (next 30 nucleotides). (C) Generation time measures end-to-end inference cost for sequence recovery on 30,000 input sequences of length 6,144 base pairs with generation of the next 50 base pairs. (D) Variant effect prediction performance is evaluated on ClinVar using AUPRC.

**Figure 4:**
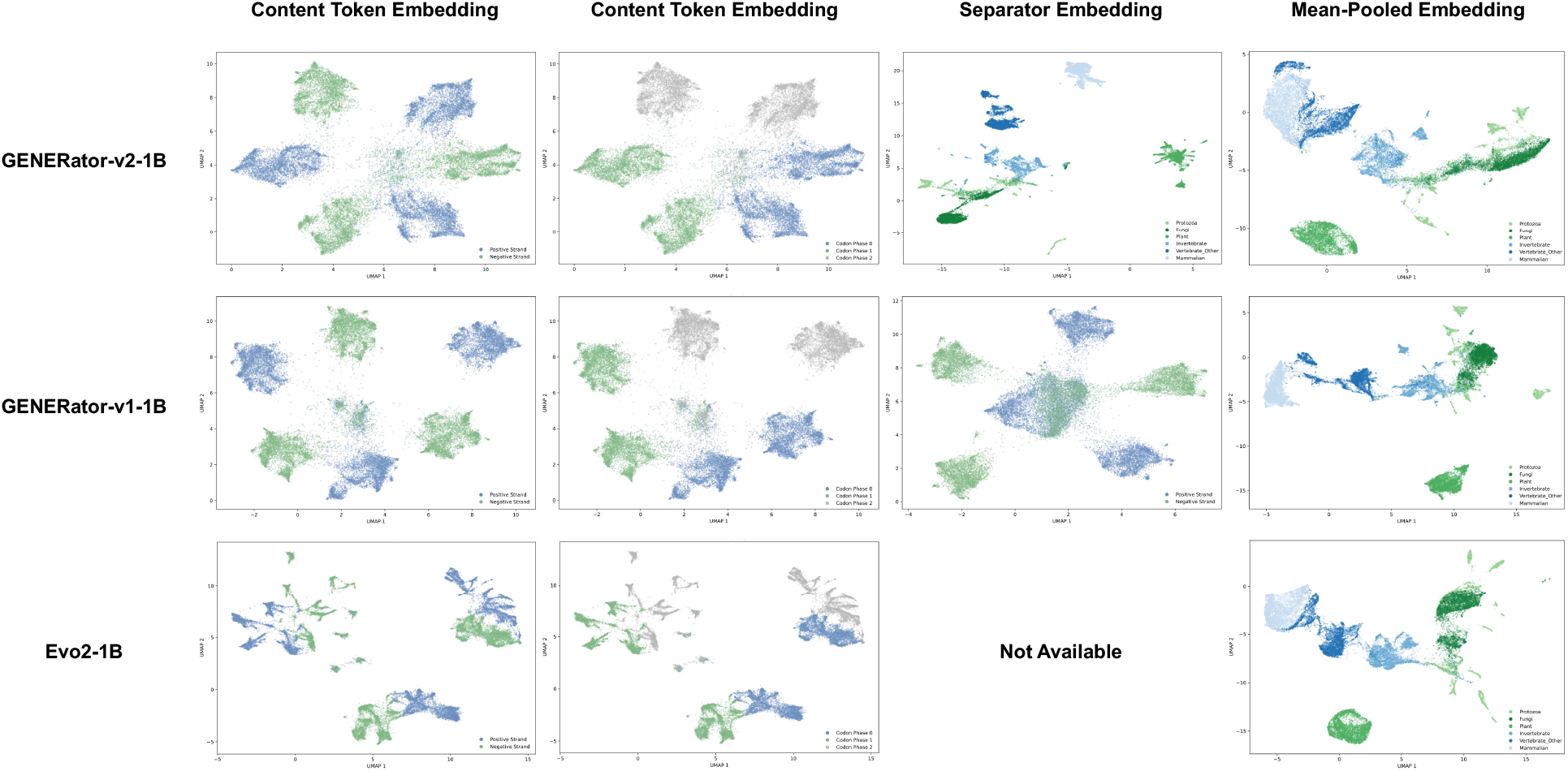
Comparison of sequence embedding structures learned by GENERator-v2, GENERator-v1, and Evo2 under different embedding extraction strategies. Rows correspond to models, and columns correspond to embedding types. *Content token embeddings* (first two columns) are obtained from the final hidden state of the last sequence token and visualized with respect to DNA strand orientation and codon phase. All models exhibit some sensitivity to these local sequence features. However, GENERator-v2 produces well-separated and highly symmetric cluster geometry across strands and codon phases, whereas GENERator-v1 shows qualitatively similar but less regular structure. Evo2 exhibits only limited separation, with noticeably weaker organization and reduced symmetry. *Separator embeddings* (third column) are obtained by appending a separator token <s> and extracting its final hidden state. Under Genome Compression Pretraining (GCP), GENERator-v2 learns an effective summary representation: separator embeddings collapse strand and codon-phase distinctions and instead cluster sequences by species, reflecting gene- or region-level information induced by the next-gene prediction objective. In contrast, GENERator-v1 does not exhibit this behavior; its separator embeddings largely retain content-like properties and remain dominated by local sequence features. Evo2 does not employ a separator or end-of-sequence token and is therefore not shown in this column. *Mean-pooled embeddings* (fourth column), obtained by averaging token embeddings across the sequence, capture coarse species-level structure in all models. However, separation is weaker than that achieved by the separator embeddings of GENERator-v2.

#### Results on prokaryotic genomes

On prokaryotic datasets, the relative performance of GENERator-v2 and Evo2 varies across bacterial clades. GENERator-v2 substantially outperforms Evo2 on groups such as *Bacillota, Epsilon-proteobacteria*, and *Gammaproteobacteria*, whereas Evo2 shows stronger performance on *Alphaproteobacteria* and *Bacteroidia*. When aggregated across all prokaryotic groups, the overall performance of the two models is comparable, and both substantially outperform Evo (Figure 3B). Notably, Evo exhibits accuracy close to random guessing (0.25) under this protocol, indicating limitations specific to the Evo model that may reflect differences in training objective and model design.

#### Generation efficiency

We further report generation time under the same evaluation setting. On a benchmark consisting of 30,000 input sequences of length 6,144 base pairs, with generation of the next 50 base pairs (corresponding to 9 predicted 6-mer tokens for GENERator), GENERator models are orders of magnitude faster than Evo-based models. Specifically, GENERator-v2-1B completes the task in 0.37 hours and GENERator-v2-3B in 0.62 hours, compared to 7.21 hours for Evo2-1B and 27.12 hours for Evo2-7B. Under the same setting, Evo-7B requires 674.52 hours while delivering the poorest recovery accuracy (Figure 3C).

#### Context length and long-context behavior

The choice of a 6,144 bp input length is not tailored to favor GENERator. This length is substantially shorter than the maximum context supported by both GENERator and Evo2, and was selected primarily to control computational cost, particularly for Evo2, whose inference cost increases rapidly with input length.

We additionally evaluated sequence recovery under shorter and longer input lengths, with detailed results reported in the Appendix. Consistent with prior observations, Evo2 exhibits limited improvement in recovery accuracy once the input length exceeds approximately 1,024 base pairs. In contrast, GENERator continues to benefit from longer input contexts. For example, when evaluated with an input length of 6,144 tokens (corresponding to 36,864 base pairs), GENERator-3B achieves further improvements in recovery accuracy. Under this configuration, its inference time is comparable to that of Evo2-7B evaluated at 6,144 base pairs, while surpassing Evo2-7B in recovery accuracy on several taxonomic groups, including *vertebrate (other), invertebrate*, and *plant* (Figure S1).

Accordingly, the results reported here should be interpreted as fair comparisons under matched input length and controlled computational cost, rather than as an upper bound on the long-context capability of GENERator.

### 5.2 Variant Effect Prediction

Sequence recovery provides a convenient training-free measure of intrinsic generative accuracy, but it does not by itself guarantee that the predicted probabilities capture biologically meaningful constraints. In particular, high recovery can arise from memorizing frequent local patterns rather than learning biologically meaningful probability structure. To further assess whether model predictions reflect genuine biological constraints, we evaluate *variant effect prediction* (VEP), a task that directly probes the calibration and interpretability of nucleotide-level probabilities.

#### Problem formulation

Given a reference sequence *S* = (*s*_1_, *s*_2_, …, *s*_*i*_, …, *s*_*L*_) and a single-nucleotide variant at position *i* with reference allele *R* and alternative allele *A*, we define the VEP score as the log-likelihood ratio

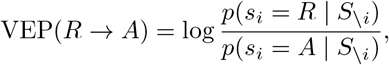

where *S \*_*i*_ denotes the sequence context excluding position *i*. Positive scores indicate preference for the reference allele and suggest stronger evolutionary constraint, while negative scores indicate preference for the alternative allele.

#### From token probabilities to nucleotide probabilities

For models using *k*-mer tokenization, direct comparison of token-level probabilities is ill-defined for single-nucleotide variants, as it involves contrasting values in a 4^*k*^-way classification space that entangle multiple nucleotide positions. We therefore compute nucleotide-level probabilities via marginalization over the *k*-mer distribution.

Let *V* = *B*^*k*^ denote the *k*-mer vocabulary. For a token covering position *i* at relative index *j ∈ {*1, …, *k}*, the probability of nucleotide *X ∈ B* at position *i* is defined as

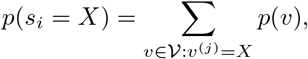

where *p*(*v*) is the model-assigned probability of *k*-mer *v* under the fixed tokenization phase used at inference, and *v*^(*j*)^ denotes the *j*-th nucleotide of *v*.

This operation mirrors the marginalization used in Factorized Nucleotide Supervision. During training, FNS applies marginalization across all *k* nucleotide positions to recover nucleotide-level supervision, whereas in variant effect prediction the same principle is applied at inference time to a single position of interest. Accordingly, VEP can be viewed as a single-position instantiation of the same nucleotide-level modeling framework. This makes variant effect prediction a natural downstream evaluation of FNS.

Because this marginalization procedure does not require architectural modification, it can be applied to all *k*-mer–based models, including GENERator-v1 and the Nucleotide Transformer series. However, GENERator-v2 is particularly well aligned with this procedure, as it is explicitly trained with FNS to optimize nucleotide-level likelihoods induced from *k*-mer logits rather than relying on token-level proxies.

#### Evaluation protocol

We evaluate VEP performance on the ClinVar dataset using the benchmark setup of GPN-MSA [5]. Inference strategies are adapted to each model class: masked language models place the variant at the sequence center with masking, while autoregressive models place the variant at the sequence end and perform next-token prediction. For all *k*-mer–based models, nucleotide-level marginalization is applied. Updated results for the NT series reflect this improved inference procedure, yielding a substantial performance gain for NT-multi-2.5B, with AUPRC increasing from 0.535 to 0.630 under nucleotide-level marginalization.

#### Results

As shown in Figure 3D, GENERator-v2 shows a clear improvement over GENERator-v1 in variant effect prediction, as measured by AUPRC. At comparable parameter scales, GENERator-v2-1B outperforms Evo2-1B, while GENERator-v2-3B matches the performance of Evo2-7B despite using approximately half the number of parameters. Other self-supervised DNA language models, including the NT series and HyenaDNA, exhibit substantially lower performance.

Alignment-based methods such as GPN-MSA [5] and CADD [32] achieve higher absolute accuracy by leveraging explicit evolutionary information from multiple sequence alignments. However, these methods are computationally expensive and largely restricted to well-studied organisms. By contrast, GENERator provides a fully alignment-free alternative that remains competitive in accuracy while being applicable across diverse species, particularly for non-model organisms.

#### Context length and computational trade-offs

To evaluate variant effect prediction under representative operating regimes, we do not enforce identical input lengths across models. GENERator-v2 is evaluated using 16,384 input tokens (approximately 98k base pairs), while Evo2 is evaluated under the configuration reported in the Evo2 paper, namely 8,192 input tokens (8,192 base pairs). According to the Evo2 authors, increasing the context length beyond this range does not lead to further improvements in variant effect prediction, a behavior that is also consistent with our observations from the sequence recovery task.

Under this protocol, GENERator-v2-3B and Evo2-7B require nearly identical inference time on the ClinVar benchmark (13.02 hours vs. 13.37 hours) and achieve the same AUPRC (0.942), despite GENERator-v2-3B using approximately half the number of parameters. At the 1B scale, GENERator-v2-1B requires less than twice the inference time of Evo2-1B (6.50 hours vs. 3.46 hours), while achieving a substantially higher AUPRC (0.934 vs. 0.906).

When evaluated with a shorter input context of 8,192 base pairs, GENERator-v2 achieves substantially faster inference while maintaining strong predictive performance. Specifically, GENERator-v2-1B completes inference in 0.36 hours with an AUPRC of 0.913, and GENERator-v2-3B requires 0.68 hours with an AUPRC of 0.928. These results demonstrate that GENERator-v2 enables a flexible and practically useful accuracy–efficiency trade-off for variant effect prediction. Longer contexts can be used to maximize performance when computational resources permit, while shorter contexts provide a highly cost-effective alternative that retains competitive accuracy.

### 5.3 Embedding Representations

We next investigate the effect of Genome Compression Pretraining (GCP) on representation learning by analyzing the structure of sequence embeddings. All evaluations in this subsection are training-free and probe the intrinsic properties of representations learned during pre-training.

#### Embedding definitions

We consider three types of sequence embeddings extracted from a pre-trained model. The *content token embedding* is obtained from the final hidden state of the last token in the input sequence. This is a common choice for autoregressive models, as under causal attention the last token is the only position whose representation can attend to the full input context. The *separator embedding* is obtained by appending a special separator token <s> to the end of the sequence and extracting the final hidden state of this token. The *mean-pooled embedding*, computed by averaging final-layer hidden states across all tokens in the sequence.

#### Evaluation dataset

To evaluate embedding structure, we construct a balanced dataset spanning six major eukaryotic taxonomic groups: *protozoa, fungi, plant, invertebrate, vertebrate (other)*, and *mammalian*. For each group, we randomly sample 5,000 genomic segments of length 8,192 base pairs from RefSeq genomes, yielding 30,000 sequences in total. These segments are sampled from random genomic locations without homology alignment. To control biological context, we require that the rightmost end of each segment lies within a coding sequence (CDS), ensuring a well-defined codon structure.

#### Visualization protocol

We project embeddings into two dimensions using UMAP [25] and visualize the resulting structure under different colorings: (i) species type, (ii) DNA strand orientation, and (iii) codon phase, defined as the alignment offset between the 6-mer tokenization window and the canonical codon boundary (phase 0: codon-aligned; phase 1/2: frame-shifted).

#### Results on GENERator-v2

The three embedding types reveal complementary properties. Content token embeddings strongly encode local sequence structure: they form well-separated clusters corresponding to DNA strand and codon phase, while species-level information remains diffuse within each cluster. Notably, the resulting cluster geometry exhibits a high degree of symmetry. Clusters associated with different strands and codon phases are arranged in a visually balanced and regular manner, with comparable separation between corresponding pairs.

This symmetry is expected from a biological and statistical perspective. DNA strand orientation is intrinsically symmetric under reverse complementation, and codon phase follows a natural three-periodic structure. In an idealized representation that faithfully captures local genomic structure, these factors should therefore be encoded in a symmetric fashion, with approximately uniform distances between equivalent strand- and phase-specific states.

Separator embeddings exhibit qualitatively different behavior. They collapse strand and codon-phase distinctions and instead form clear clusters aligned with species taxonomy. This divergence is consistent with the training dynamics induced by GCP. During pre-training, tokens preceding the separator are optimized primarily for local next-token prediction, encouraging embeddings that capture strand orientation and codon periodicity. Once the model encounters <s>, the prediction objective shifts: the separator embedding must summarize the preceding region in order to predict the next neighboring functional region. We refer to this behavior as *next-gene prediction*. As a result, <s> functions not only as a separator but also as an explicit summary token, yielding a high-quality gene-level embedding. This behavior provides a representation-level signature of functional in-context learning, where information from multiple upstream functional units is integrated to inform downstream predictions.

Mean-pooled embeddings provide a simpler global baseline. By aggregating token-level representations across the sequence, mean pooling captures coarse global information and yields clustering aligned with species taxonomy. However, the resulting separation is noticeably weaker and less well defined than that achieved by separator embeddings, indicating that simple pooling lacks the discriminative power of a learned summary representation.

#### Comparison with GENERator-v1

GENERator-v1 exhibits broadly similar behavior to GENERator-v2 in terms of content token embeddings and mean-pooled sequence embeddings, indicating that both models learn comparable representations of local sequence structure and coarse taxonomic signal. However, a clear divergence emerges for separator embeddings. In GENERator-v1, the separator embedding fails to function as a meaningful summary representation and largely retains the characteristics of content token embeddings, in some cases exhibiting even weaker structure.

This behavior is expected, as GENERator-v1 does not employ GCP and therefore lacks an explicit next-gene prediction objective that would encourage the separator token to aggregate information across functional regions. As a result, the separator embedding in GENERator-v1 does not acquire the distinct semantic role observed in GENERator-v2. This behavior provides a representation-level signature of functional in-context learning induced by GCP.

Beyond the separator token, more subtle differences are also apparent. Content token embeddings in GENERator-v1 exhibit less regular and less symmetric cluster geometry than those in GENERator-v2, and mean-pooled embeddings show slightly weaker separation across taxonomic groups. Together, these observations suggest that the improvements in GENERator-v2 reflect not only the introduction of functional-region compression, but also more effective supervision that enhances representation quality at both local and global levels.

#### Comparison with Evo2

We further include Evo2-1B as a representative single-nucleotide–tokenized baseline. Unlike GENERator, Evo2 does not employ an explicit separator or end-of-sequence token in its vocabulary. Accordingly, we do not construct a separator embedding for Evo2. Instead, we compare content token embeddings and mean-pooled sequence embeddings across models.

Mean-pooled embeddings from Evo2-1B exhibit clustering behavior broadly similar to those of GENERator-v1 and GENERator-v2, forming coherent groups aligned with species taxonomy. This suggests that, at the level of global sequence summaries, Evo2 learns representations that capture coarse phylogenetic structure.

In contrast, Evo2 content token embeddings show substantially weaker structure. While Evo2-1B exhibits a limited ability to distinguish DNA strand orientation and codon phase, the resulting clusters are noticeably less well separated and less symmetric than those observed in GENERator models. In addition, extracting Evo2 embeddings incurs substantially higher inference cost than GENERator at comparable sequence lengths, reflecting the computational overhead of single-nucleotide tokenization.

Notably, we also observe a layer-dependent effect in Evo2 representations. Evo2-1B consists of 24 StripedHyena2 layers; however, representations extracted from the final three layers exhibit extremely large activation magnitudes (on the order of 10^10^) and do not display meaningful clustering structure. In contrast, embeddings from earlier layers remain stable and progressively improve in structure with depth up to this point. This behavior is consistent with observations reported in the Evo2 study [7], and we therefore restrict our analysis to the 21st layer, which is the deepest layer that remains numerically stable and interpretable.

#### Discussion

These results establish a desirable representational organization for genomic foundation models trained for functional in-context learning. Without ever observing explicit labels for strand orientation, codon structure, or species identity—and despite the absence of homology alignment—GENERator-v2 learns representations that separate local syntactic features from higher-level biological semantics. Content token embeddings predominantly encode local sequence properties such as strand orientation and codon phase, while separator embeddings act as effective summary representations that cluster genomic segments according to species taxonomy, even when the segments originate from distant genomic loci. This separation of roles indicates that the model captures genuine genomic regularities rather than memorizing positional or dataset-specific artifacts.

The geometry of the learned representations provides an additional internal consistency check. Although GENERator-v2 operates on a 6-mer tokenizer, clustering with respect to reading-frame alignment follows the natural three-nucleotide periodicity of codons, corresponding to phases 0, 1, and 2, rather than exhibiting an artificial 6-periodic structure. Moreover, strand- and phase-specific clusters display a high degree of symmetry. This behavior is biologically expected, reflecting the symmetry of double-stranded DNA and the equivalence of different codon frame offsets. The emergence of this structure suggests that, when coupled with appropriate supervision and training objectives, *k*-mer tokenization does not obscure biologically meaningful syntax.

Taken together, these comparisons highlight that single-nucleotide tokenization is not inherently superior to *k*-mer tokenization in terms of representation quality. At matched model scale, GENERator models consistently exhibit more structured and better organized embeddings—particularly at the level of content token representations—despite operating on a coarser tokenization. Moreover, these representational advantages are achieved with substantially lower computational cost at inference time. Together, these findings suggest that pre-training objective and data construction play a more critical role than token granularity alone, and that well-aligned supervision can enable *k*-mer–based models to capture fine-grained genomic structure at least as effectively as single-nucleotide–based alternatives.

### 5.4 Benchmark Evaluations

In addition to training-free evaluations, we assess downstream performance under task-specific fine-tuning to verify that the improvements introduced by FNS and GCP transfer to standard supervised genomic tasks. We evaluate fine-tuning performance on the revised Nucleotide Transformer (NT) tasks [9], which comprise a diverse collection of genomic classification problems and address several limitations of the original NT benchmark. A number of alternative benchmark suites, including Genomic Benchmarks [13], GUE [42], and BEND [24], have been used in prior work. Rather than exhaustively evaluating across all available benchmarks, we focus on the revised NT tasks as a representative and well-established fine-tuning setting.

#### Fine-tuning protocol

We do not include Evo2 in fine-tuning experiments because end-to-end fine-tuning at scale is computationally prohibitive in practice. Under training-free inference, Evo2-1B is already more than an order of magnitude slower than GENERator-3B in our evaluations (Figure 3C), and the computational gap would be expected to increase further under full fine-tuning. This practical constraint likely explains why, to the best of our knowledge, no prior work has reported end-to-end fine-tuning results for Evo2 on established genomic benchmarks.

For the same reason, we apply this criterion consistently to our own models and restrict fine-tuning experiments to GENERator-v2-1B, excluding the 3B variant. While larger models are expected to achieve stronger absolute performance, such gains would primarily reflect scaling effects rather than qualitative differences in training objective or data construction. All fine-tuning experiments follow a consistent protocol with hyperparameter search, a linear classification head, and 10-fold cross-validation; full details are provided in the Appendix.

#### Results

Across the revised NT benchmark, GENERator-v2 consistently outperforms GENERator-v1 on the majority of tasks and achieves near-equivalent performance on the remainder (Table 2). These results indicate that the improvements introduced by FNS and GCP persist under supervised task adaptation and are not limited to training-free evaluation settings.

**Table 2:** Evaluation of the revised Nucleotide Transformer tasks. The reported values represent the Matthews correlation coefficient (MCC) averaged over 10-fold cross-validation, with the standard error in parentheses.

|  | Enformer<br>(252M) | DNABERT-2<br>(117M) | HyenaDNA<br>(55M) | NT<br>(2.5B) | NT-v2<br>(500M) | Caduceus-Ph<br>(8M) | Caduceus-PS<br>(8M) | GROVER<br>(87M) | GENERator-v1<br>(1.2B) | GENERator-v2<br>(1.2B) |
| --- | --- | --- | --- | --- | --- | --- | --- | --- | --- | --- |
| H2AFZ | 0.522 (0.019) | 0.490 (0.013) | 0.455 (0.015) | 0.503 (0.010) | 0.524 (0.008) | 0.417 (0.016) | 0.501 (0.013) | 0.509 (0.013) | 0.529 (0.009) | **0.533 (0.005)** |
| H3K27ac | 0.520 (0.015) | 0.491 (0.010) | 0.423 (0.017) | 0.481 (0.020) | 0.488 (0.013) | 0.464 (0.018) | 0.464 (0.022) | 0.489 (0.023) | **0.546 (0.015)** | 0.544 (0.012) |
| H3K27me3 | 0.552 (0.007) | 0.599 (0.010) | 0.541 (0.018) | 0.593 (0.016) | 0.610 (0.006) | 0.547 (0.010) | 0.561 (0.036) | 0.600 (0.008) | 0.619 (0.008) | **0.623 (0.003)** |
| H3K36me3 | 0.567 (0.017) | 0.637 (0.007) | 0.543 (0.010) | 0.635 (0.016) | 0.633 (0.015) | 0.543 (0.009) | 0.602 (0.008) | 0.585 (0.008) | 0.650 (0.006) | **0.659 (0.006)** |
| H3K4me1 | 0.504 (0.021) | 0.490 (0.008) | 0.430 (0.014) | 0.481 (0.012) | 0.490 (0.017) | 0.411 (0.012) | 0.434 (0.030) | 0.468 (0.011) | 0.504 (0.010) | **0.518 (0.006)** |
| H3K4me2 | **0.626 (0.015)** | 0.558 (0.013) | 0.521 (0.024) | 0.552 (0.022) | 0.552 (0.013) | 0.480 (0.013) | 0.526 (0.035) | 0.558 (0.012) | 0.607 (0.010) | 0.616 (0.011) |
| H3K4me3 | 0.635 (0.019) | 0.646 (0.008) | 0.596 (0.015) | 0.618 (0.015) | 0.627 (0.020) | 0.588 (0.020) | 0.611 (0.015) | 0.634 (0.011) | 0.653 (0.008) | **0.657 (0.014)** |
| H3K9ac | **0.593 (0.020)** | 0.564 (0.013) | 0.484 (0.022) | 0.527 (0.017) | 0.551 (0.016) | 0.514 (0.014) | 0.518 (0.018) | 0.531 (0.014) | 0.570 (0.017) | 0.569 (0.008) |
| H3K9me3 | 0.453 (0.016) | 0.443 (0.025) | 0.375 (0.026) | 0.447 (0.018) | 0.467 (0.044) | 0.435 (0.019) | 0.455 (0.019) | 0.441 (0.017) | 0.509 (0.013) | **0.517 (0.018)** |
| H4K20me1 | 0.606 (0.016) | 0.655 (0.011) | 0.580 (0.009) | 0.650 (0.014) | 0.654 (0.011) | 0.572 (0.012) | 0.590 (0.020) | 0.634 (0.006) | **0.670 (0.006)** | 0.669 (0.004) |
| Enhancer | **0.614 (0.010)** | 0.517 (0.011) | 0.475 (0.006) | 0.527 (0.012) | 0.575 (0.023) | 0.480 (0.008) | 0.490 (0.009) | 0.519 (0.009) | 0.594 (0.013) | 0.580 (0.007) |
| Enhancer type | **0.573 (0.013)** | 0.476 (0.009) | 0.441 (0.010) | 0.484 (0.012) | 0.541 (0.013) | 0.461 (0.009) | 0.459 (0.011) | 0.481 (0.009) | 0.547 (0.017) | 0.546 (0.010) |
| Promoter all | 0.745 (0.012) | 0.754 (0.009) | 0.693 (0.016) | 0.761 (0.009) | 0.780 (0.012) | 0.707 (0.017) | 0.722 (0.014) | 0.721 (0.011) | **0.795 (0.005)** | **0.795 (0.005)** |
| Promoter non-TATA | 0.763 (0.012) | 0.769 (0.009) | 0.723 (0.013) | 0.773 (0.010) | 0.785 (0.009) | 0.740 (0.012) | 0.746 (0.009) | 0.739 (0.018) | **0.801 (0.005)** | **0.801 (0.006)** |
| Promoter TATA | 0.793 (0.026) | 0.784 (0.036) | 0.648 (0.044) | 0.944 (0.016) | 0.919 (0.028) | 0.868 (0.023) | 0.853 (0.034) | 0.891 (0.041) | 0.950 (0.009) | **0.956 (0.013)** |
| Splice acceptor | 0.749 (0.007) | 0.837 (0.006) | 0.815 (0.049) | 0.958 (0.003) | **0.965 (0.004)** | 0.906 (0.015) | 0.939 (0.012) | 0.812 (0.012) | 0.964 (0.003) | 0.964 (0.003) |
| Splice site all | 0.739 (0.011) | 0.855 (0.005) | 0.854 (0.053) | 0.964 (0.003) | 0.968 (0.003) | 0.941 (0.006) | 0.942 (0.012) | 0.849 (0.015) | 0.966 (0.003) | **0.970 (0.002)** |
| Splice donor | 0.780 (0.007) | 0.861 (0.004) | 0.943 (0.024) | 0.970 (0.002) | 0.976 (0.003) | 0.944 (0.026) | 0.964 (0.010) | 0.842 (0.009) | 0.977 (0.002) | **0.978 (0.002)** |

#### Discussion

Fine-tuning benchmarks remain a widely used and historically important evaluation paradigm for genomic foundation models, and have played a central role in assessing task-specific performance. However, as models continue to scale, the practical limitations of fine-tuning become increasingly pronounced. End-to-end fine-tuning is computationally expensive for large models, and fine-tuning pipelines often involve many task-specific design choices, leading to substantial variability across implementations and reduced reproducibility. As a result, fine-tuning benchmarks can be difficult to compare reliably across studies.

Moreover, fine-tuning updates model parameters for individual tasks, which can partially obscure or overwrite representational differences learned during pre-training. This makes fine-tuning less sensitive to advances in intrinsic model quality, particularly when the goal is to compare pre-training objectives or data construction strategies.

A similar shift has been observed in the development of large language models, where early progress was primarily evaluated through fine-tuning benchmarks (e.g., in the BERT era [39, 36]), while more recent work increasingly emphasizes zero-shot and training-free evaluations as models scale [14, 8]. Motivated by this perspective, our experiments demonstrate that training-free evaluations provide signals that are largely consistent with fine-tuning performance, while offering improved reproducibility, lower computational cost, and clearer cross-model comparison. We therefore advocate broader use of training-free evaluation protocols alongside fine-tuning benchmarks to support more robust and transparent progress in genomic language modeling.

## 6 Discussion

Taken together, our results motivate a re-examination of genomic foundation model design through the lens of training objective, data construction, and evaluation philosophy. In particular, this work focuses on two persistent bottlenecks in genomic language modeling: the mismatch between coarse tokenization and nucleotide-level biology, and the extreme sparsity of functional signal in eukaryotic genomes.

Factorized Nucleotide Supervision (FNS) addresses the supervision axis by reconciling efficient *k*-mer tokenization with single-nucleotide resolution. Despite operating in a 4^*k*^-way token space, the resulting training objective remains equivalent to optimizing nucleotide-level likelihoods. Genome Compression Pretraining (GCP) addresses the data construction axis by reshaping the training distribution, allowing models to allocate capacity to biologically informative regions and to learn higher-level structure through next-gene prediction, a representation-level manifestation of functional in-context learning. The combination of these techniques yields GENERator-v2 models that consistently improve over GENERator-v1 across comprehensive evaluations while maintaining favorable computational efficiency.

### Limitations

Despite these advances, several limitations remain. First, GCP relies on the availability of high-quality genomic annotations. While raw genomic sequence is increasingly easy to obtain, comprehensive and accurate annotations remain scarce, unevenly distributed across species, and costly to produce. This dependency constrains the applicability of GCP in poorly annotated genomes and highlights a gap between sequence availability and functional understanding.

Second, GENERator inherits intrinsic limitations of autoregressive models. While causal attention is well suited for generative and probabilistic tasks, it is suboptimal for dense, position-wise annotation. Annotating positions near the beginning of a sequence requires access to both upstream and downstream context, which causal attention cannot provide.

### Future directions

These limitations naturally motivate two complementary research directions. The first concerns annotation itself. To reduce reliance on curated annotations, one promising direction is the development of alignment-free gene annotation models. In parallel work, we explore this direction through GENERanno [17], a companion model that employs bidirectional attention to access full sequence context when predicting functional labels. Such models offer a pathway toward scalable, automated annotation that can in turn support annotation-driven training strategies such as GCP.

The second direction concerns extending the principles of efficient tokenization and nucleotide-level supervision beyond autoregressive modeling. In this work, we demonstrate that *k*-mer tokenization combined with FNS yields both high efficiency and single-nucleotide fidelity for causal language models. In principle, similar ideas could be applied to masked language models, enabling efficient and nucleotide-resolved DNA annotation at scale. However, the MLM setting introduces additional challenges, most notably the need to mask and supervise only subsets of nucleotides within a *k*-mer token. Designing principled masking and loss formulations in this regime remains an active area of our ongoing work.

### Outlook

These results suggest a modular view of genomic foundation models. Efficient generative modeling, accurate nucleotide-level probability estimation, and dense functional annotation need not be addressed by a single architecture. Instead, specialized models—such as GENERator for generation and probabilistic reasoning, and annotation-oriented models with bidirectional context—can be combined into a coherent ecosystem, in which improvements in one component reinforce the others. This perspective provides a practical and scalable path toward genomic foundation models that more faithfully reflect the structure and constraints of genomic data.

Finally, as part of this broader vision, we are committed to advancing genomic language modeling through open and reproducible research. To promote transparency, reproducibility, and broad adoption, all model checkpoints, pretraining data artifacts, and scripts for downstream analyses associated with this work are released publicly at https://huggingface.co/GenerTeam. We invite the research community to build upon these resources and to participate in collaborative efforts aimed at accelerating progress in genomic language modeling and functional genomics.

## A Factorized Nucleotide Supervision (FNS)

This appendix provides formal definitions, theoretical results, and algorithmic details supporting Section 2. In particular, we show that although GENERator is trained with a *k*-mer tokenizer, the Factorized Nucleotide Supervision (FNS) objective is equivalent to predicting the next *k* single-nucleotide tokens under a shared conditioning state.

### A.1 Setup and Notation

Let ℬ = {*A, C, G, T*} denote the nucleotide alphabet and let 𝒱 = ℬ^*k*^ denote the *k*-mer vocabulary with size *V* = |𝒱| = 4^*k*^. At autoregressive step *t*, the model produces logits **z**_*t*_ ∈ ℝ^V^, inducing a categorical distribution over *k*-mers

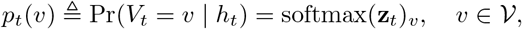

where *h*_*t*_ denotes the conditioning state (e.g., the decoder hidden state determined by *x*_*<t*_).

Each *k*-mer token *v* ∈ 𝒱 corresponds to an ordered tuple of nucleotides *v* = (*v*^(1)^, …, *v*^(*k*)^) with *v*^(*j*)^ *∈ B*. For nucleotide position *j* ∈ {1, …, *k*} and base *b* ∈ ℬ, we define the induced nucleotide marginal

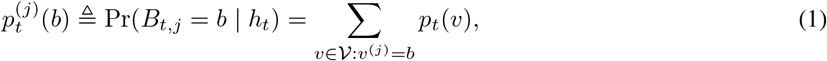

which is a valid categorical distribution over ℬ.

Given a ground-truth *k*-mer target 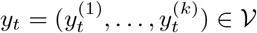, the FNS per-step loss is defined as

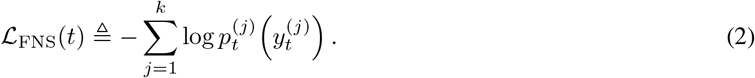

### A.2 Formal Results

#### Definition 1

(Blockwise 1-mer predictor). Fix a step *t* with conditioning state *h*_*t*_. A *blockwise 1-mer predictor* outputs *k* categorical distributions 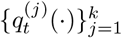 over *B* and defines a joint distribution over the next nucleotide block 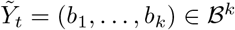 via conditional independence:

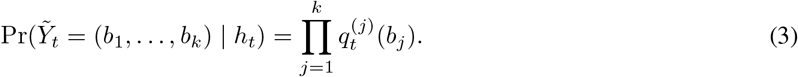

This corresponds to predicting *k* consecutive 1-mer tokens in parallel, conditioned on the same *h*_*t*_.

#### Theorem 2

(FNS is equivalent to blockwise 1-mer maximum likelihood). *Let* 𝒱 = ℬ^*k*^ *be the k-mer vocabulary and let p*_*t*_(*v*) = softmax(**z**_*t*_)_*v*_ *be the model distribution over k-mers at step t. Define induced nucleotide marginals* 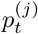 (*·*) *by Eq*. (1). *Consider a blockwise 1-mer predictor with per-position distributions set to*

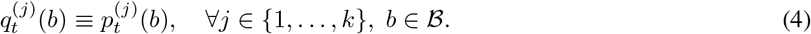

*Then for any ground-truth k-mer target* 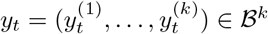, *the FNS loss equals the negative log-likelihood of predicting the next k 1-mer tokens under the blockwise 1-mer predictor:*

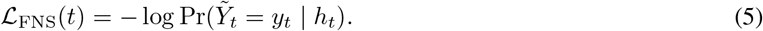

*Proof*. By Definition 1 and Eq. (4),

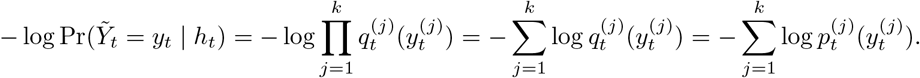

The right-hand side is exactly the FNS loss by Eq. (2), proving Eq. (5).

**Implication for nucleotide resolution**. Theorem 2 shows that, although the model predicts a single *k*-mer token at each step, the learning objective supervises each of the *k* nucleotide positions through an explicit 4-way categorical likelihood. Therefore, *k*-mer tokenization does not degrade single-nucleotide resolution under FNS; instead, it provides a computationally efficient reparameterization of blockwise single-nucleotide prediction.

#### Corollary 3

(SNP likelihood ratio under blockwise 1-mer equivalence). *Let y, y*^*′*^ ∈ ℬ^*k*^ *differ only at position j. Under the blockwise 1-mer interpretation in Theorem 2*,

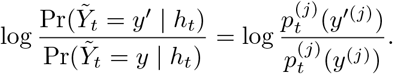

*Proof*. By Eq. (3),

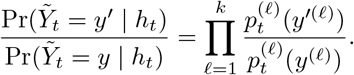

All factors cancel except *𝓁* = *j* since *y*^*′*(*𝓁*)^ = *y*^(*𝓁*)^ for *𝓁≠ j*.

### A.3 6-mer Tokenization under FNS vs. Single-Nucleotide Tokenization

Under FNS, next *k*-mer prediction can be interpreted as predicting the next *k* single-nucleotide tokens in parallel under a conditional-independence assumption given the shared conditioning state (Theorem 2). This factorization is an explicit approximation at the level of the training objective: it restores nucleotide-level supervision while retaining the computational benefits of coarse *k*-mer tokenization.

Beyond conditional independence, *k*-mer tokenization also changes the *granularity of attention*. Compared to single-nucleotide tokenization, a 6-mer token effectively bundles multiple adjacent nucleotides into a single query/key/value (Q/K/V) unit, i.e., attention is computed over *blocks* rather than individual bases. This blockwise representation can reduce the model’s ability to attend to base-level microstructure *within* a block at a single attention step, which may be disadvantageous for very short sequences (e.g., 30 bp) where fine-grained local interactions dominate and the number of available tokens is already small.

However, for long genomic sequences, this trade-off is typically negligible and can be beneficial in practice. When sequence length is large, the dominant challenge is allocating limited attention and model capacity across long-range dependencies and multiple functional units. Blockwise attention reduces the number of tokens by a factor of *k*, enabling substantially larger effective genomic span and more stable optimization under a fixed compute budget. Empirically, this perspective is consistent with our sequence recovery results, where 6-mer models trained with FNS maintain strong nucleotide-level fidelity while achieving large gains in efficiency and benefiting from longer contexts (Figure S1).

### A.4 Algorithmic Details

Algorithm 1 summarizes the computation of the FNS loss for *k*-mer tokenized DNA sequences. The algorithm marginalizes token-level probabilities into nucleotide-level probabilities and applies a token-level fallback for special tokens (e.g., padding or separators), yielding a numerically stable and fully differentiable implementation of Eq. (2).

## B Genome Compression Pretraining (GCP)

This appendix provides formal definitions, theoretical analyses, and algorithmic details supporting Section 3. We formalize the construction of compressed genomic sequences, analyze the effect of tokenization-shift augmentation under Factorized Nucleotide Supervision (FNS), and examine representation stability in comparison to sparse attention mechanisms.

### B.1 Setup and Notation

**Nucleotide alphabet and sequences**. Let ℬ = *{A, C, G, T}* denote the nucleotide alphabet. A DNA sequence is a finite string *S* = (*b*_1_, …, *b*_*n*_) ∈ ℬ^*n*^.

**Reference genomes and annotations**. A reference genome *G* consists of chromosomes {*C*}. Each chromosome *C* has an associated DNA sequence seq(*C*) and length |*C*|. Functional annotations are represented as intervals

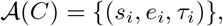

where 0 *≤ s*_*i*_ *< e*_*i*_ *≤* |*C*| and *τ*_*i*_ denotes the annotation type.

#### Algorithm 1

Factorized Nucleotide Supervision (FNS) Loss

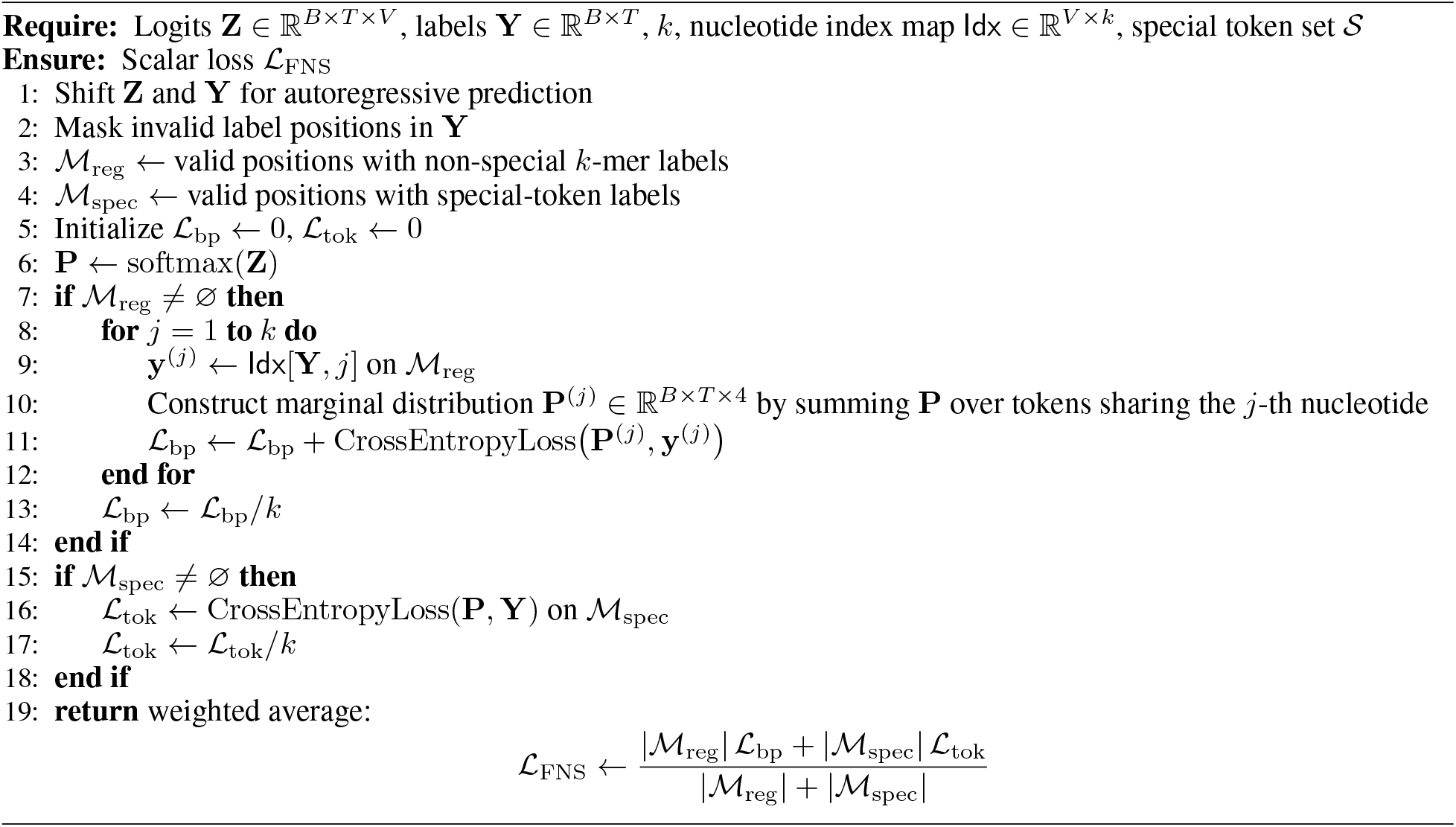

#### Non-overlapping *k*-mer tokenization

Let *k* denote the *k*-mer size. A non-overlapping tokenization with offset *δ* ∈ {0, …, *k −* 1} maps *S* = (*b*_1_, …, *b*_*n*_) to

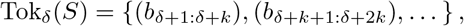

discarding incomplete *k*-mers at the boundaries.

#### Model and hidden states

We consider an autoregressive decoder parameterized by *θ*. Given previous tokens **x**_*<t*_, the model produces a hidden state *h*_*t*_ ∈ ℝ^*d*^and logits 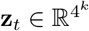 over the *k*-mer vocabulary.

#### Separator token

<s> denotes a special token used to mark boundaries between functional regions. Separator tokens are included in the input but masked from the language modeling loss.

### B.2 Formal Results

Non-overlapping *k*-mer tokenization is inherently phase-sensitive: shifting the tokenization offset *δ* by one nucleotide yields a completely different sequence of token IDs. When combined with long-context training, this raises a natural concern: does the choice of tokenization phase affect which nucleotide positions receive supervision during training?

We show that under Factorized Nucleotide Supervision (FNS), this concern does not arise. Specifically, when all *k* tokenization offsets are exhaustively enumerated during training, the resulting supervision signal is complete and consistent at the nucleotide level, up to boundary effects that can be eliminated by padding.

#### B.2.1 Nucleotide-Level Loss Induced by FNS

Let *S* = (*b*_1_, …, *b*_*n*_) ∈ ℬ^*n*^ be a DNA sequence. For a fixed offset *δ* ∈ {0, …, *k*− 1}, let Tok_*δ*_(*S*) produce *m*_*δ*_ non-overlapping *k*-mer tokens. Under FNS, each token contributes *k* nucleotide-level negative log-likelihood terms. We define the total nucleotide-level loss induced by offset *δ* as

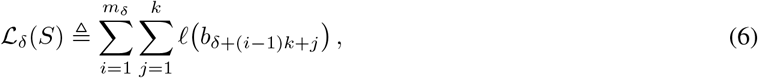

where *𝓁*(*b*_*t*_) denotes the negative log-probability assigned to nucleotide *b*_*t*_ under FNS, and indices outside {1, …, *n*} are omitted.

##### Algorithm 2

Genome Compression Pretraining (GCP)

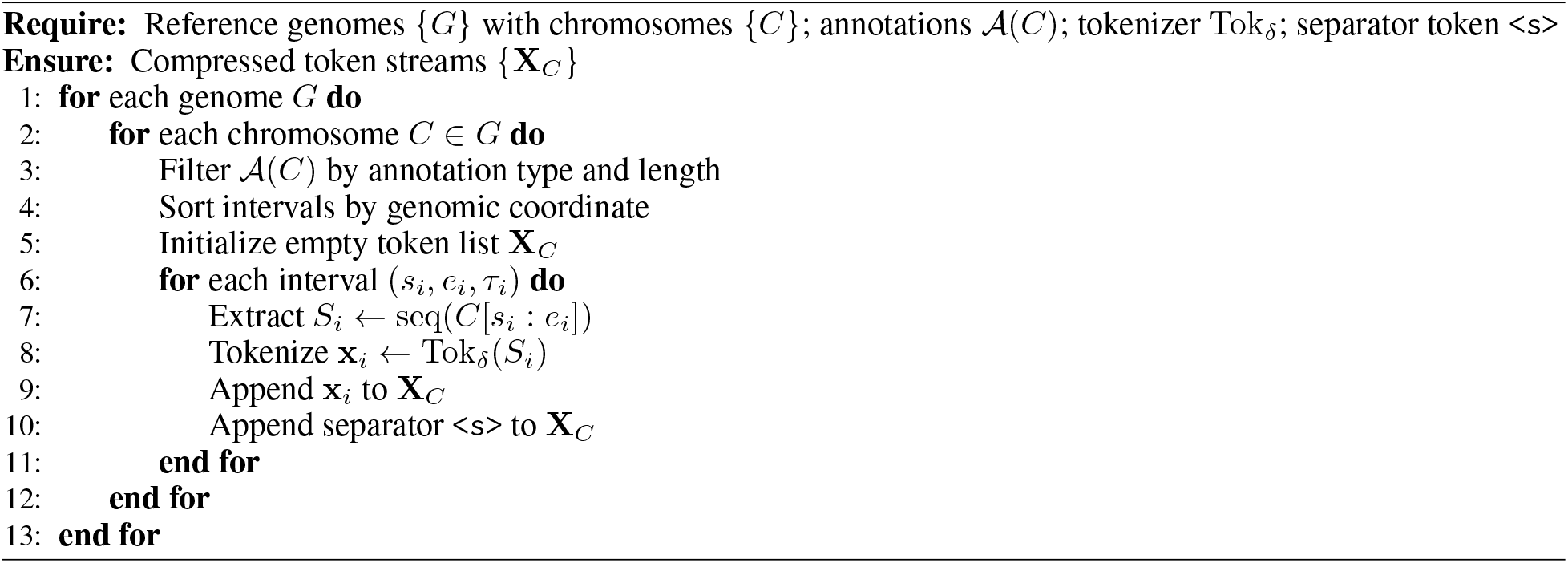

#### B.2.2 Phase-Invariant Nucleotide Supervision under Tokenization Shifts

Because different offsets *δ* cover different subsets of nucleotide positions, the per-offset losses *L*_*δ*_(*S*) are not identical. However, in training protocols that exhaustively cycle through all *k* offsets (e.g., one offset per epoch), the relevant quantity is the *combined* nucleotide-level supervision across offsets.

##### Proposition 4

(Tokenization-phase invariance under FNS). *Let S* = (*b*_1_, …, *b*_*n*_) ∈ ℬ^*n*^ *and k* ≥ 1. *For each offset δ ∈ {0, …, *k* − 1}, define L*_*δ*_(*S*) *as in Eq. (6). Then the total nucleotide-level loss aggregated across all offsets satisfies*

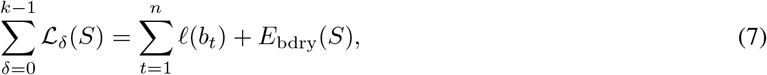

*where E*_bdry_(*S*) *depends only on nucleotides within k*− 1 *positions of the sequence boundaries. Moreover, if S is padded on both sides with at least k*− 1 *dummy nucleotides that are masked from the loss, then E*_bdry_(*S*) = 0 *and Eq. (7) holds exactly*.

*Proof*. Fix an offset *δ* and define the set of nucleotide positions covered by complete *k*-mers under *δ* as

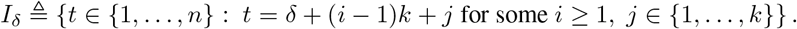

By definition, 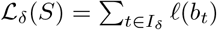.Summing over offsets yields

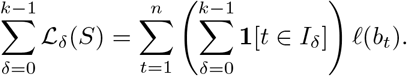

For any interior position *t* ∈ {*k*, …, *n− k*} + 1, there exists exactly one offset *δ ≡ t −*1 (mod *k*) such that *t ∈ I*_*δ*_, hence the inner sum equals 1. Boundary positions may be excluded for some offsets due to incomplete *k*-mers; these discrepancies are collected into *E*_bdry_(*S*). Padding with masked dummy nucleotides ensures every original position belongs to exactly one *I*_*δ*_, yielding *E*_bdry_(*S*) = 0.

### B.3 Algorithmic Details

Algorithm 2 describes the construction of compressed training sequences used in GCP. Training windows of fixed length *L* tokens are sampled from **X**_*C*_ using random offsets or a fixed stride, and loss terms corresponding to separator tokens are masked.

## C Benchmark Evaluations

We select the revised NT tasks for fine-tuning performance evaluation. Following GENERator-v1 [41], we evaluate GENERator-v2 using 10-fold cross-validation and use the Matthews correlation coefficient (MCC) for main metric.

To align with GENERator-v2’s 6-mer tokenization, we truncate sequences to the closest multiple of 6 during training and evaluation. We place a separator token (<s>) at both the start and end of the truncated sequence, and use the final-layer hidden state of the terminal <s> token as the sequence embedding, which is then passed through a linear classification head for downstream prediction. We conduct extensive hyperparameter searches, exploring learning rates in {1 × 10^−5^, 2 × 10^−5^, 5 × 10^−5^} and batch sizes in {32, 64, 128, 256}. We maintain pre-training optimizer settings (*β*_1_ = 0.9, *β*_2_ = 0.95, weight decay = 0.1) and implement a reduce-on-plateau learning rate scheduler with 100 steps of linear warm-up and early stopping with patience=5. Optimal hyperparameters identified through grid search are documented in Table S1.

## D Supplementary Figures and Tables

**Figure S1:**
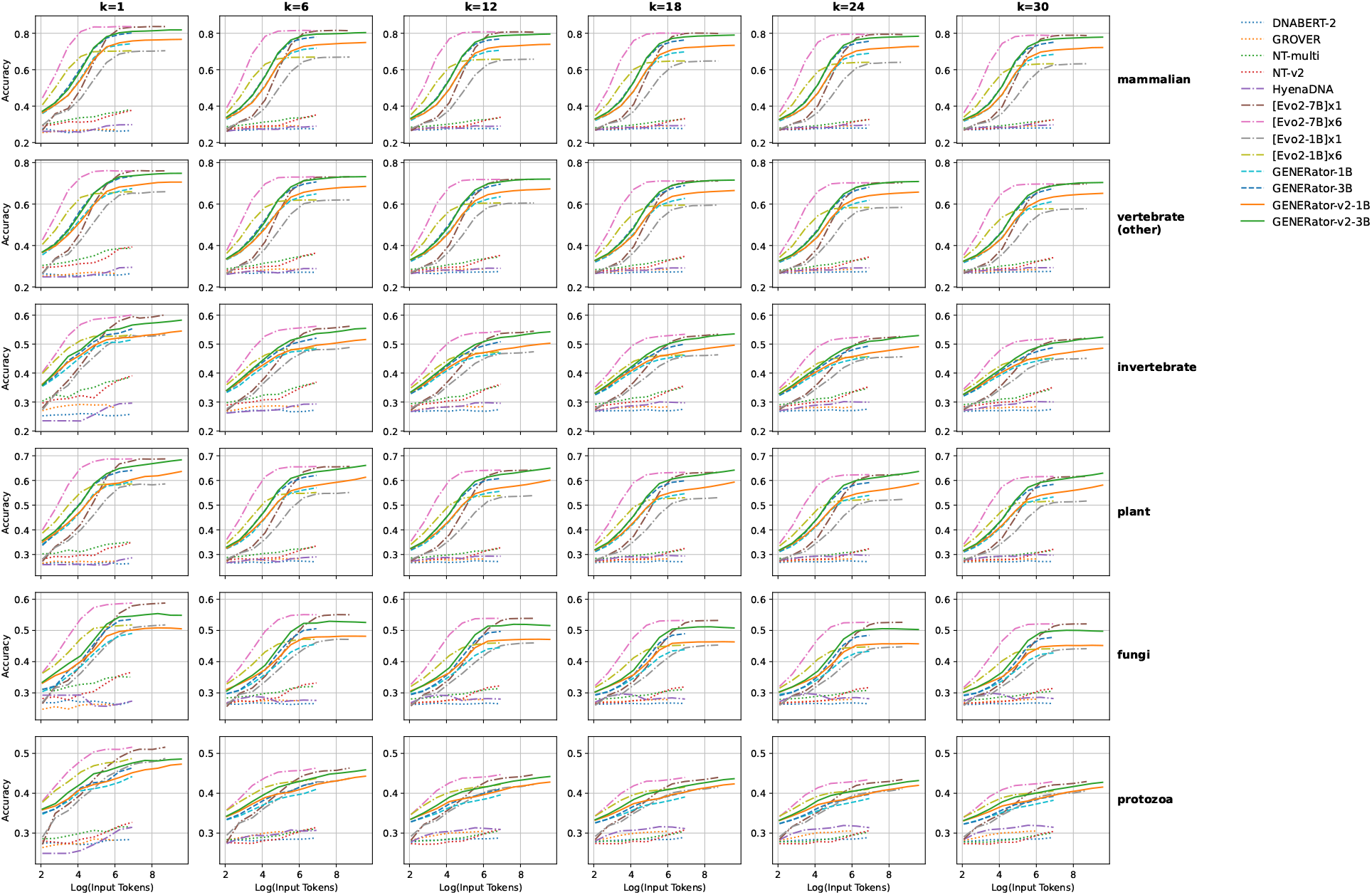
Sequence recovery accuracy as a function of input length across genomic foundation models. Each panel reports recovery accuracy versus the number of input tokens (log-scaled) for a specific taxonomic group and prediction length. Curves correspond to different model families, including MLM-based genomic foundation models (e.g., DNABERT-2, GROVER, and the NT series) and autoregressive models such as HyenaDNA, Evo2, and the GENERator family. MLM-based models consistently exhibit poor performance, as expected due to their lack of autoregressive generative capability. HyenaDNA, while autoregressive, is severely limited by its small parameter count (55M), resulting in substantially lower accuracy despite its long nominal context length. For Evo2, we report two configurations: [Evo2]×1, which uses the same token count as other models and reflects comparable computational efficiency, and [Evo2]×6, which matches the nucleotide count of GENERator by using six times more tokens at the expense of an order-of-magnitude increase in inference cost. Although [Evo2]×6 shows an initial advantage at very short input lengths, this advantage diminishes rapidly, and [Evo2]×1 (1,024 tokens) and [Evo2]×6 (6,144 tokens) converge to nearly identical accuracy, indicating limited utilization of extended context. In contrast, both GENERator and GENERator-v2 exhibit strong parameter efficiency and sustained gains with increasing input length. GENERator-v2 consistently improves over GENERator across taxonomic groups, demonstrating more effective utilization of longer contexts and translating additional input into meaningful accuracy gains.

**Table S1:** Hyperparameter settings for the revised Nucleotide Transformer tasks.

|  | GENERator-v2 |  |
| --- | --- | --- |
|  | LR | BS |
| H2AFZ | $1e^{-5}$ | 32 |
| H3K27ac | $2e^{-5}$ | 32 |
| H3K27me3 | $1e^{-5}$ | 32 |
| H3K36me3 | $1e^{-5}$ | 128 |
| H3K4me1 | $2e^{-5}$ | 64 |
| H3K4me2 | $2e^{-5}$ | 64 |
| H3K4me3 | $2e^{-5}$ | 64 |
| H3K9ac | $1e^{-5}$ | 32 |
| H3K9me3 | $2e^{-5}$ | 32 |
| H4K20me1 | $1e^{-5}$ | 64 |
| Enhancer | $2e^{-5}$ | 32 |
| Enhancer types | $2e^{-5}$ | 32 |
| Promoter all | $2e^{-5}$ | 256 |
| Promoter non-TATA | $1e^{-5}$ | 32 |
| Promoter TATA | $1e^{-5}$ | 32 |
| Splice acceptor | $2e^{-5}$ | 32 |
| Splice site all | $2e^{-5}$ | 32 |
| Splice site donors | $2e^{-5}$ | 64 |

